# Genetic parameters of colostrum and calf serum antibodies in Swedish dairy cattle

**DOI:** 10.1101/2021.08.08.455591

**Authors:** Juan Cordero-Solorzano, Dirk-Jan de Koning, Madeleine Tråvén, Therese de Haan, Mathilde Jouffroy, Andrea Larsson, Aline Myrthe, Joop A.J. Arts, Henk K. Parmentier, Henk Bovenhuis, Jonas Johansson Wensman

## Abstract

**Background:** Colostrum with sufficient IgG content is essential for the newborn calf, as it provides passive immunity which substantially affects the probability of survival during rearing. Failure of passive transfer occurs when a calf does not absorb enough antibodies from the colostrum as indicated by less than 10 g/L of IgG in calf serum. Besides delayed access to colostrum, this can be due to low IgG production of the mother or poor IgG absorption by the calf. The aim of this study was to estimate the genetic background of antibody levels and indicator traits for antibodies in colostrum and calf serum, and their correlation with milk production and health.

**Results:** Colostrum data were available from 1340 cows with at least one calving and calf serum data were available from 886 calves from these cows. Antibody concentrations were estimated using refractometry (digital Brix refractometer for colostrum and optical refractometer for serum) as indicator traits and established using ELISAs to determine total IgG and natural antibodies [NAb] of various antibody isotypes in colostrum and serum. Colostrum traits had heritabilities ranging from 0.16 to 0.31 with repeatabilities from 0.21 to 0.55. Brix had positive genetic correlations with all colostrum antibody traits including total IgG (0.68). Calf serum antibody concentrations had heritabilities ranging from 0.25 to 0.59, with a significant maternal effect accounting for 17 to 27% of the variance. When calves later in life produced their first lactation, lactation-average somatic cell score was found to be negatively correlated with NAb in calf serum.

**Conclusions:** Our results suggest that antibody levels in colostrum and calf serum can be increased by means of selection.

## Background

At birth, calves are highly dependent on the absorption of maternal antibodies from the colostrum to acquire passive humoral immunity and local protection of the digestive tract. If not enough antibodies are transferred, failure of passive transfer (**FPT**) occurs, defined as < 10 g/L of IgG or < 5.5 g/dL of serum total protein (**STP**) present in calf serum at 24 hours after birth [1]. Calves with FPT have twice the risk of morbidity and death at a young age compared to calves with proper serum IgG (**S- IgG**) levels [2]. Estimates of the prevalence of FPT in Swedish dairy herds range from 14% [3] to 60% [4]. Assessing colostrum quality is an important factor in preventing FPT. It should contain at least 50 g/L of IgG or a Brix value larger than 22%, to be considered of sufficient quality to transfer passive immunity to the calf [5].

There is a large variation in colostrum quality between cows, even between animals of the same farm and breed [6]. Some of this variation can be explained by environmental factors, but there is an important genetic component in colostrum antibody content. A previous study estimated a heritability of 0.4 (standard error = 0.3) for colostrum IgG [7]. More recently, a study by Soufleri et al. (2019) found a heritability of 0.27 (0.09) for Brix percentage in colostrum of Holstein cows.

Despite its importance for calf health, very few studies have focused on the genetics associated with calf antibody uptake from colostrum. It has been observed that even when the time of first meal, volume of first meal, colostrum antibody concentration and other variables have been accounted for, there is still significant unexplained variation left in calf antibody uptake [9]. Part of this variation can be due to genetics. Gilbert et al. (1988) estimated a heritability of 0.56 (0.25) for calf serum IgG1 in 36 hour-old calves using a paternal half-sib analysis accounting for the dam’s colostral IgG1 concentration while Burton et al. (1989) estimated a heritability of 0.18 (0.25) for IgG in calves 24 to 36 h old using a paternal half-sib analysis. More recently, Martin et al. (2021) using 366 Charolais calves estimated a heritability of 0.36 (0.18) for calf serum IgG at 24 to 48h after birth.

The most frequent diseases during rearing are diarrhea (scours) and pneumonia, with diarrhea being the most prevalent for calves younger than 30 days of age and pneumonia for weaners over 30 days of age [12]. These diseases are caused by a variety of bacteria, viruses and parasites [13]. Farms with high prevalence of calf diarrhea and pneumonia face economic losses from higher mortality and increased costs from purchasing replacement heifers to make up for the lost ones [14].

Natural antibodies (**NAb**) are immunoglobulins produced by a subset of B cells (in human and mice) known as B-1 cells, as a part of the innate immune system, without any antigenic stimulation [15]. These antibodies likely provide first defense towards a variety of infectious diseases, tumors and metabolic disorders [16]. In dairy cattle NAb levels have been associated with risk of mastitis [17], length of productive life [18], lameness [19] and postpartum uterine health [20]. In poultry, serum NAb levels have been linked to survival [21] and *E. coli* resistance [22].

The aim of this study was to measure genetic components of antibody levels and potential indicator traits in colostrum and calf serum, and their correlation with milk production and health.

## Methods

### Samples

The samples were collected at three experimental farms in Sweden: The Swedish Livestock Research Center Lövsta (Uppsala), Röbäcksdalen forskningsstation SITES (Umeå) and Nötcenter Viken (Falköping) from January 2015 to April 2017. An overview of the traits and number of samples by farm and breed can be found in Table 1.

**Table 1.**
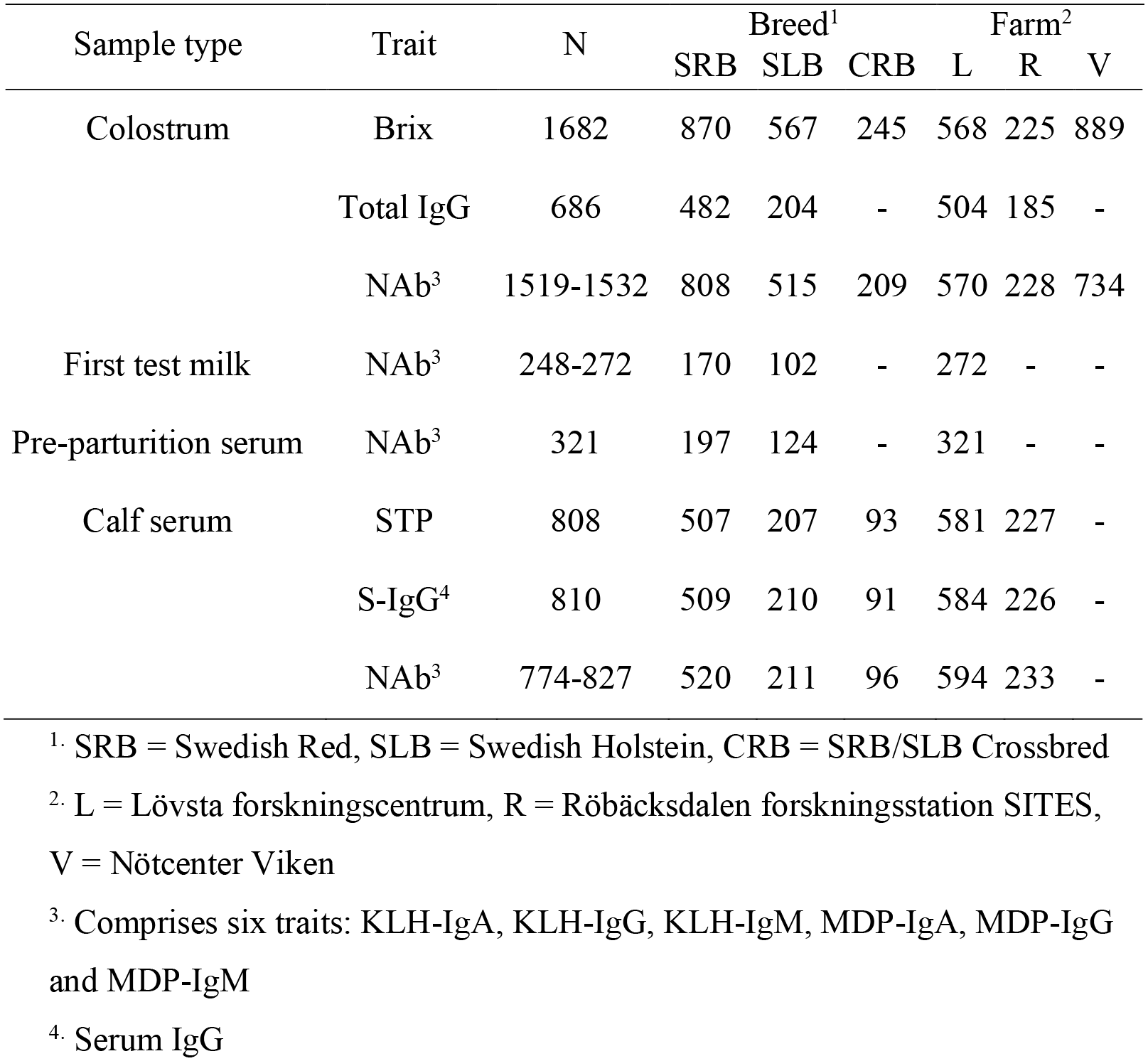
Overview of traits and number of samples by breed and farm.

| Sample type | Trait | N | Breed<sup>1</sup> |  |  | Farm<sup>2</sup> |  |  |
| --- | --- | --- | --- | --- | --- | --- | --- | --- |
|  |  |  | SRB | SLB | CRB | L | R | V |
| Colostrum | Brix | 1682 | 870 | 567 | 245 | 568 | 225 | 889 |
|  | Total IgG | 686 | 482 | 204 | - | 504 | 185 | - |
|  | NAb<sup>3</sup> | 1519-1532 | 808 | 515 | 209 | 570 | 228 | 734 |
| First test milk | NAb<sup>3</sup> | 248-272 | 170 | 102 | - | 272 | - | - |
| Pre-parturition serum | NAb<sup>3</sup> | 321 | 197 | 124 | - | 321 | - | - |
| Calf serum | STP | 808 | 507 | 207 | 93 | 581 | 227 | - |
|  | S-IgG<sup>4</sup> | 810 | 509 | 210 | 91 | 584 | 226 | - |
|  | NAb<sup>3</sup> | 774-827 | 520 | 211 | 96 | 594 | 233 | - |

<sup>1</sup>. SRB = Swedish Red, SLB = Swedish Holstein, CRB = SRB/SLB Crossbred
<sup>2</sup>. L = Lövsta forskningscentrum, R = Röbäcksdalen forskningsstation SITES, V = Nötcenter Viken
<sup>3</sup>. Comprises six traits: KLH-IgA, KLH-IgG, KLH-IgM, MDP-IgA, MDP-IgG and MDP-IgM
<sup>4</sup>. Serum IgG

### Cows

In total 1,340 cows were sampled, out of which 682 were Swedish Red (**SRB**), 460 Swedish Holstein (**SLB**) and 198 SLB and SRB crossbreds (**CRB**). Parities ranged from 1 to 6 with 90% of the animals in parity 1 to 3. During the sampling period 504 cows calved twice and 29 calved three times.

Sampling of first milking colostrum was carried out by staff at the farms, recording the time between birth and sampling for a total of 1,711 individual samples. Additionally, pre-parturition serum and first test milk samples were taken only from cows at Lövsta; 330 serum samples from January 2015 to June 2016, 23 to 1 days before calving and 290 milk samples between February 2015 to March 2016 at 6 to 24 days in milk.

Milk production information was provided by Växa Sverige for 1,283 cows, comprising 305-day milk yield (kg), fat and protein percentage and lactation-average somatic cell score (**LASCS**) from lactations matching the corresponding colostrum sample. LASCS values were calculated from somatic cell count (**SCC**) measurements as described by Wijga et al. (2012).

### Calves

A total of 886 calves were sampled at Lövsta and Röbäcksdalen, borne by cows with a matching colostrum sample. However, 59 calves could not be linked to a pedigree record and were excluded from the study. The remaining 827 calves were 520 SRB, 211 SLB and 96 CRB.

Neonates were separated from the dam as soon as possible after birth. At Röbäcksdalen, all calves were fed with nipple bottle while in Lövsta, they were predominantly fed by nipple bottle and those too weak to suckle were fed by esophageal tube. All calves were given first milking colostrum as the first meal.

Sampling of blood was carried out by staff at the farms. Blood was drawn using a vacutainer system (BD Biosciences, NJ, USA) for collection of serum, in tubes without additives, and anti-coagulated blood in tubes with K_3_-ethylenediaminetetraacetic acid (**EDTA**). Anti-coagulated blood was used for DNA extraction and genotyping.

Calves were sampled at the age of 1 to 12 days, but 93% of the samples were taken before 8 days. For each individual calf, birth weight, volume of first meal, time of first meal after birth, and colostrum donor of first meal, was recorded.

Disease information and treatments during the first three months of age were recorded for 233 calves born in Röbäcksdalen during the sampling period. Additional information on veterinary treatments of animals from Lövsta and Röbäcksdalen at 8 to 36 months of age was made available by Växa Sverige. In both cases physical trauma reports were excluded.

First parity milk production information was provided by Växa Sverige for 253 of the animals that started the study as calves, comprising 305-day cumulative milk yield (kg), fat and protein percentage and LASCS.

### Phenotypes

#### Brix

Colostrum samples were analyzed using a digital refractometer Atago 3810 PAL-1 (Atago, Tokyo, Japan) to measure Brix percentage, which approximates total solids (**TS**) percentage, as an estimate for IgG content. A total of 1,682 individual samples were analyzed. Each sample was measured three times to average a sample value.

#### Serum total protein (STP)

An optical refractometer AO Veterinary Refractometer 10436 (AO Scientific Instruments, NY, USA) was used to measure TS in calf serum samples as an estimate of STP (g/dL). A total of 808 samples were analyzed. Each sample was measured three times to average a sample value.

#### Total IgG in colostrum

Colostrum IgG was measured only in samples with a matching calf serum sample using a commercial ELISA (Bovine IgG ELISA Kit E-10G, Immunology Consultants Laboratory Inc, OR, USA) according to the manufacturer’s instructions. Standards and samples were run in duplicates and a four-parameter logistic regression was used to calculate the IgG concentration (g/L). Colostrum (686 samples) were diluted at 1:400,000 but samples with high values outside the standard reference curve limits were retested at 1:800,000.

#### Total IgG in calf serum

Serum IgG concentration was determined using an ELISA kit (Bovine IgG ELISA Quantitation Set E10-118, Bethyl Laboratories Inc, TX, USA) according to the manufacturer’s instructions. Standards and samples were run in duplicate and a logit regression was used to calculate IgG concentration (g/L). Sera (810 samples) were diluted at 1:112,000; however, samples with values outside the standard reference curve limits were retested at 1:224,000 or 1:16,000 for higher or lower values, respectively.

To avoid confusion with the NAb traits, these IgG measurements will be referred to as “colostrum total IgG” or simply “total IgG” for colostrum and “serum IgG” or “S-IgG” for calf serum.

#### Natural Antibodies

Titers for NAbs were measured in all sample types for cows (colostrum, first test milk and pre-parturition serum) and calves (serum). Optical density (**OD**) of muramyl dipeptide (**MDP**) and keyhole limpet hemocyanin (**KLH**)-binding immunoglobulins of the isotypes IgM, IgA, and IgG were measured by an indirect two-step ELISA as outlined by Ploegaert et al. (2010).

Colostrum and serum samples were prediluted at 1:10, whereas milk samples at 1:5, with phosphate-buffered saline containing 0.05% Tween 20 (**PBST** pH 7.2, dilution buffer). Flat-bottomed, 96-well medium binding plates were coated with 100 μl/well of 2 μg/ml of KLH (H8283, Sigma-Aldrich), or MDP (A9519, Sigma-Aldrich) respectively, in carbonate buffer (5.3 g/L Na2CO3 and 4.2 g/L NaHCO3, pH 9.6). After incubation overnight at 4°C, plates were washed with tap water containing Tween 20 and blocked with 100 μl/well of 5% normal chicken serum in PBST for one hour at room temperature.

Prediluted samples were further diluted in the antigen-coated plates with dilution buffer to 1:40, 1:160, 1:640, and 1:2,560 test dilutions for colostrum and serum and 1:10, 1:20, 1:40, and 1:80 for milk. One unrelated colostrum sample was chosen as standard positive to be consistently used in all assays. Duplicates of this standard positive were stepwise 1:2 diluted with dilution buffer (8 serial dilutions from 1:20 until 1:2,560) and pipetted into the antigen-coated plates. The plates were incubated for one hour and a half at room temperature.

After washing, MDP or KLH plates were incubated with 100 μl/well of either 1:40,000-diluted rabbit-anti-bovine IgM labelled with horseradish peroxidase (**HRP**) (A10-100P, Bethyl Laboratories Inc), 1:40,000-diluted sheep-anti-bovine IgG HRP (A10-118P, Bethyl Laboratories Inc) or 1:20,000- diluted rabbit-anti-bovine IgA HRP (A10-108P, Bethyl Laboratories Inc) for one hour and a half at room temperature. After washing, 100 µL substrate buffer (containing water, 10% tetramethylbenzidine (**TMB**) buffer [15 g/L sodium acetate and 1.43 g/L urea hydrogen peroxide; pH 5.5], and 1 % tetramethylbenzidine [8 g/L TMB in DMSO]) was added and incubated for approximately ten minutes at room temperature. The reaction was stopped with 50 μL of 1.25M H_2_SO_4_. OD was measured with a Multiskan Go (Thermo Scientific, MA, USA) at OD 450 nm.

Antibody titers were calculated as described by Frankena (1987) [cited by de Koning et al. (2015)]. Optical densities of the duplicate standard positive samples were averaged for each plate. Logit values of OD per plate were calculated using:

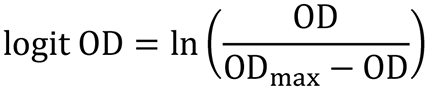

where OD is the OD of a well, and OD_max_ is the maximum averaged OD of the duplicate standard positive samples. The last positive well (**lpw**) of the averaged duplicate standard positive was set to the sixth dilution for colostrum and serum and to the seventh for milk. Titers of each sample per plate were calculated using:

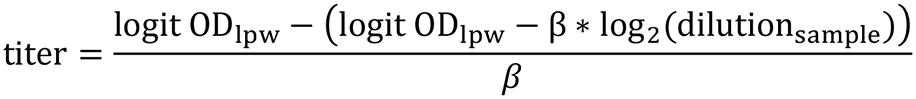

where logit OD_lpw_ is the estimated logit OD at the lpw calculated with the estimated linear regression function using the log_2_-dilution value of that well, logit OD_sample_ is the logit OD calculated of the OD closest to 50% of OD_max_ for a sample of an individual (OD_sample_), β is a regression coefficient of the logit OD against the respective log_2_-dilution values of the averaged duplicate standard positive samples, and log_2_ dilution_sample_ is the log_2_-dilution value at which OD_sample_ occurred, as described by de Koning et al. (2015).

Six traits were generated for each sample type from three isotypes measured for two antigens: KLH- IgA, KLH-IgG, KLH-IgM, MDP-IgA, MDP-IgG and MDP-IgM. A total of 1,510 to 1,532 samples (depending on the trait) were analyzed for each colostrum trait, 248 to 272 for first test milk, 321 for pre-parturition cow serum and 774 to 827 for calf serum. A constant was added to each trait to make all the values larger than zero.

### Statistical analysis

Variance components for genetic effects, repeated measurements (colostrum traits) and maternal effects (calf serum) were estimated with animal models using ASReml 4.1 [27]. The pedigree used for all animals contained 29,048 records (20 generations) and was made available by Växa Sverige.

#### Models for cows

For colostrum traits, the following repeatability model was used:

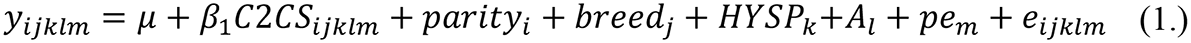

where y is the observation of the trait; μ is the overall mean of the trait; C2CS_ijklm_ is a covariate describing the effect of colostrum sampling time after calving in hours; parity_i_ represents the fixed effect of four parity classes (1, 2, 3 and 4 or more); breed_j_ is the fixed effect of breed (SLB, SRB or CRB); HYSP_k_ describes the fixed effect of Herd-Year-Season of calving and sample storage Plate number; A_l_ the random additive genetic effect assumed to be distributed as 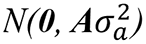, where **A** is the additive genetic relationships matrix from the pedigree and 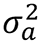 is the additive genetic variance; pe_m_ is the random permanent environment effect assumed to be distributed as 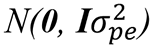, where **I** is the identity matrix and 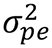 is the permanent environment effect variance; e_ijklm_ is the random residual effect assumed to be distributed as 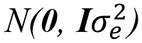 where **I** is the identity matrix and 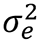is the residual variance.

For first test milk and pre-parturition cow serum NAb traits, from cows at Lövsta experimental farm the following animal model was used:

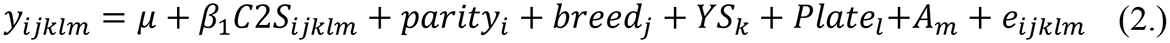

where y is the NAb titer of milk or serum; μ is the overall mean of the trait; C2S_ijklm_ is a covariate describing the effect of sampling time in days before (serum) or after (milk) calving; parity_i_ represents the fixed effect of four parity classes (1, 2, 3 and 4 or more); breed_j_ is the fixed effect of breed (SLB, SRB or CRB); YS_k_ describes the fixed effect of Year-Season of calving; Plate_l_ is the fixed effect of sample storage Plate number; A_m_ the random additive genetic effect assumed to be distributed as 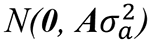, where **A** is the additive genetic relationships matrix from the pedigree and 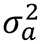 is the additive genetic variance; e_ijklm_ is the random residual effect assumed to be distributed as 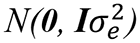 where **I** is the identity matrix and 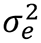 is the residual variance.

#### Models for calves

For calf serum traits, the following animal model was used:

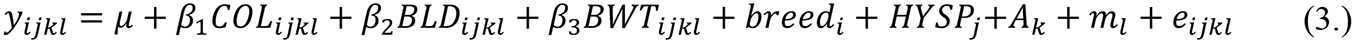

where y is the observation of the trait; μ is the overall mean; COL_ijkl_ is a covariate describing the absolute amount of antibodies received from the matching colostrum trait (Brix, colostrum IgG or NAb * volume of first meal); BLD_ijkl_ is a covariate describing the time from calving to blood sampling in days; BWT_ijkl_ is the covariate of calf birth weight in kg; breed_i_ is the fixed effect of breed (SLB, SRB or CRB); HYSP_j_ describes the fixed effect of Herd-Year-Season of calving and sample storage Plate number; A_m_ the random additive genetic effect assumed to be distributed as 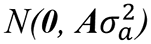, where **A** is the additive genetic relationships matrix from the pedigree and 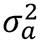 is the additive genetic variance; m_l_ is the random maternal effect assumed to be distributed as 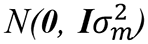, where **I** is the identity matrix and 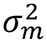 is the maternal variance; e_ijkl_ is the random residual effect assumed to be distributed as 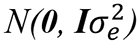, where **I** is the identity matrix and 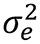 is the residual variance.

#### Heritabilities and variance proportions

Heritabilities were estimated as:

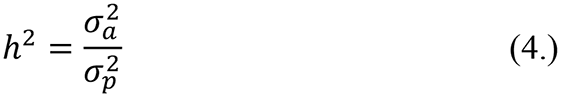

with phenotypic variance 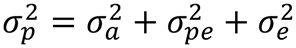 for model (1.), 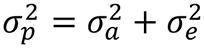 for model (2.) and 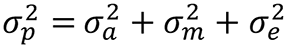 for model (3.), where 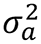 is the additive genetic variance, 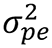 the permanent environment variance, 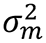 is the maternal variance and 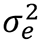 the residual variance.

Repeatability for colostrum traits was:

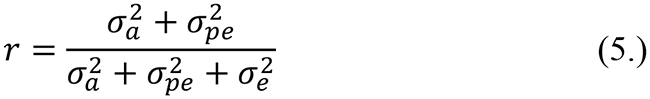

where 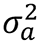 is the additive genetic variance, 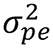 the permanent environment variance and 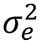 the residual variance.

Maternal contribution or maternal variance proportion for calf traits was estimated as:

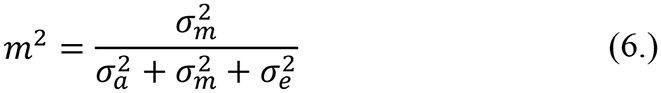

Genetic correlations were estimated using bivariate analyses between different colostrum traits (model 1) and between different calf serum traits (model 3) as follows:

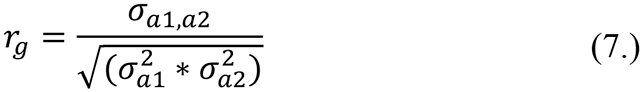

where 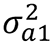 is the additive genetic variance for trait 1, 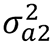 the additive genetic variance for trait 2, and *σ*_*a*1,*a*2_ the additive genetic covariance between traits 1 and 2. The same formula was applied for phenotypic correlations, substituting additive genetic variances and covariance for phenotypic ones. Genetic and phenotypic correlations were also estimated between colostrum (model 1) and calf serum traits (model 2). A similar approach was used to estimate genetic and phenotypic correlations of colostrum traits (model 1) with first test milk (model 2) and pre-parturition cow serum (model 3). Significance of correlations was checked using a log-likelihood ratio test, heritabilities and genetic correlations were considered significant if they were different from 0.

### Health and production traits analysis

#### Cows

Using available milk production data for 305 days average of milk yield, fat and protein percentage and LASCS, genetic and phenotypic correlations were estimated between these and colostrum traits. The following repeatability model was used for milk production traits:

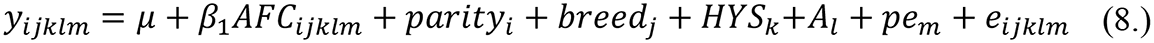

where y is the observation of the milk trait; μ is the overall mean; AFC_ijklm_ is a covariate describing the effect of age at first calving in months; parity_i_ represents the fixed effect of four parity classes (1, 2, 3 and 4 or more); breed_j_ is the fixed effect of breed (SLB, SRB or CRB); HYS_k_ describes the fixed effect of Herd-Year-Season of calving; A_l_ the random additive genetic effect assumed to be distributed as 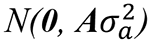, where **A** is the additive genetic relationships matrix from the pedigree and 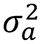 is the additive genetic variance; pe_m_ is the random permanent environment effect assumed to be distributed as 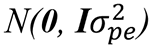, where **I** is the identity matrix and 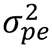 is the permanent environment effect variance; e_ijklm_ is the random residual effect assumed to be distributed as 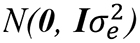, where **I** is the identity matrix and 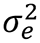 is the residual variance.

A bivariate analysis with model (1.) for colostrum traits and model (8.) for production traits was used to estimate the correlations according to formula (7.).

#### Calves

First parity milk production data from 253 of the calves was combined with 700 records of first parity milk data from the project calves’ dams. Genetic and phenotypic correlations were estimated between calf serum traits and milk production. A bivariate analysis with model (3.) for serum traits and a modified model (8.) removing the fixed effect of parity and the random effect of permanent environment, was used for production traits and age at first calving (**AFC**) as dependent variables to estimate the correlations according to formula (7.).

Disease data for Röbäcksdalen calves (0 to 3 months of age) was coded as binary (0 = healthy/untreated and 1 = disease/treated) with most of the positive cases treated with antibiotics. In total, 63 out of the 231 animals were diagnosed with at least one disease event and received treatment. A bivariate analysis with model (3.) for serum traits and a model including only Year-Season of birth as a fixed effect for health trait, was used to estimate the correlations according to formula (7.).

Health information for cattle between 8 to 36 months of age in Lövsta and Röbäcksdalen included 66 animals with reported cases out of the 301 heifer calves from our samples that were kept for milk production. Data was arranged as binary (0 = healthy and 1 = disease) regardless of the condition. A bivariate analysis with model (3.) for serum traits and a model including only Herd-Year-Season of birth as a fixed effect for health trait, was used to estimate the correlations according to formula (7.).

## Results

An overview of the traits and genetic parameters for calf serum traits is presented in Table 2, and cow colostrum, milk and serum traits in Table 3. For calf serum traits, heritability estimates were moderate to high, ranging from 0.25 to 0.59, except for STP, which was not significantly different from 0. Maternal contribution was moderate (0.17 to 0.27), except for KLH-IgA, which was not significantly different from 0. Breed was only significant for STP with an effect size of 0.27 (0.10) for SLB relative to SRB.

**Table 2.** Descriptive statistics, heritabilities (*h*^2^), maternal effects (*m*^2^), phenotypic variances 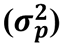, breed effects and number of samples for calf serum traits. SE in parenthesis.

| Trait<sup>1</sup> | Mean<br>(SD) | Range<sup>2</sup> | $h^2$ | $m^2$ | $\sigma_p^2$ | Breed<sup>3</sup> | | N |
| --- | --- | --- | --- | --- | --- | --- | --- | --- |
|  |  |  |  |  |  | SLB | CRB |  |
| STP (g/dL) | 6.0 (0.7) | 4.9 – 7.2 | 0.07 (0.11) | **0.22 (0.08)** | 0.44 (0.03) | **0.27 (0.10)** | **0.18 (0.09)** | 808 |
| S-IgG (g/L)<sup>4</sup> | 22.8 (12.8) | 5.6 – 48.7 | **0.25 (0.13)** | **0.27 (0.08)** | 1.28 (0.08) | 0.04 (0.24) | 0.06 (0.18) | 810 |
| KLH-IgA | 4.0 (1.3) | 1.8 – 6.2 | **0.46 (0.14)** | 0.03 (0.08) | 0.85 (0.05) | 0.06 (0.24) | -0.01 (0.16) | 827 |
| KLH-IgG | 9.3 (1.7) | 6.3 – 11.6 | **0.26 (0.14)** | **0.37 (0.08)** | 1.57 (0.10) | 0.19 (0.27) | -0.21 (0.19) | 827 |
| KLH-IgM | 6.1 (1.7) | 3.1 – 8.8 | **0.23 (0.13)** | **0.23 (0.08)** | 2.24 (0.13) | 0.04 (0.31) | -0.04 (0.23) | 827 |
| MDP-IgA | 5.8 (1.8) | 2.6 – 8.4 | **0.43 (0.16)** | **0.17 (0.08)** | 1.63 (0.10) | -0.03 (0.33) | 0.16 (0.22) | 774 |
| MDP-IgG | 6.7 (1.4) | 4.3 – 9.0 | **0.24 (0.14)** | **0.25 (0.08)** | 1.33 (0.08) | 0.14 (0.25) | -0.10 (0.18) | 821 |
| MDP-IgM | 7.8 (1.8) | 4.4 – 10.0 | **0.59 (0.16)** | **0.19 (0.08)** | 1.61 (0.10) | 0.08 (0.35) | 0.03 (0.22) | 825 |

<sup>1</sup>. Significant $h^2$ , $m^2$ and breed effects are highlighted in bold
<sup>2</sup>. Range shows 5th quantile and 95th quantile, respectively
<sup>3</sup>. Breed effects are relative to Swedish Red (SRB)
<sup>4</sup>. Variance components and effects were estimated using a square root transformation

**Table 3.** Descriptive statistics, heritabilities (*h*^2^), repeatabilities (*r*), phenotypic variances 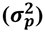, breed effects and number of samples for cow sample type and traits. SE in parenthesis.

| Trait<sup>1</sup> | Mean<br>(SD) | Range<sup>2</sup> | $h^2$ | $r$ | $\sigma_p^2$ | Breed<sup>3</sup> | | N |
| --- | --- | --- | --- | --- | --- | --- | --- | --- |
|  |  |  |  |  |  | SLB | CRB |  |
| *Colostrum* |  |  |  |  |  |  |  |  |
| Brix (%) | 21.9 (4.2) | 14.5 – 28.7 | **0.31 (0.06)** | **0.35 (0.04)** | 15.47 (0.59) | **2.02 (0.82)** | **0.86 (0.52)** | 1682 |
| Total IgG (g/L)<sup>4</sup> | 56.8 (26.9) | 19.7 – 106.6 | **0.20 (0.09)** | **0.21 (0.08)** | 2.49 (0.14) | **0.68 (0.32)** | - | 686 |
| KLH-IgA | 5.9 (1.3) | 3.8 – 8.0 | **0.26 (0.06)** | **0.45 (0.04)** | 1.37 (0.06) | **0.84 (0.23)** | **0.41 (0.16)** | 1532 |
| KLH-IgG | 8.0 (1.6) | 5.1 – 10.3 | **0.16 (0.07)** | **0.45 (0.04)** | 2.10 (0.08) | **0.94 (0.24)** | **0.50 (0.18)** | 1519 |
| KLH-IgM | 7.0 (1.3) | 5.0 – 8.7 | **0.24 (0.07)** | **0.40 (0.05)** | 1.40 (0.06) | **0.78 (0.23)** | **0.47 (0.16)** | 1532 |
| MDP-IgA | 6.4 (1.4) | 4.0 – 8.6 | **0.29 (0.07)** | **0.38 (0.05)** | 1.61 (0.07) | **0.71 (0.26)** | **0.43 (0.17)** | 1532 |
| MDP-IgG | 8.4 (1.8) | 5.5 – 11.3 | **0.24 (0.07)** | **0.55 (0.04)** | 2.84 (0.12) | **0.89 (0.33)** | **0.61 (0.22)** | 1532 |
| MDP-IgM | 9.1 (1.4) | 6.5 – 10.9 | **0.22 (0.06)** | **0.37 (0.05)** | 1.78 (0.07) | **0.80 (0.25)** | **0.60 (0.17)** | 1532 |
| *First test milk* |  |  |  |  |  |  |  |  |
| KLH-IgA | 2.1 (1.4) | 1.2 – 5.2 | 0.03 (0.17) | - | 1.53 (0.14) | **0.46 (0.18)** | - | 272 |
| KLH-IgG | 3.1 (1.0) | 1.6 – 4.9 | **0.33 (0.17)** | - | 0.95 (0.09) | **0.48 (0.26)** | - | 272 |
| KLH-IgM | 2.4 (0.9) | 1.3 – 3.8 | 0.00 (0.00) | - | 0.72 (0.06) | **0.36 (0.11)** | - | 272 |
| MDP-IgA | 2.9 (1.3) | 1.3 – 5.4 | 0.15 (0.20) | - | 1.43 (0.14) | **0.45 (0.25)** | - | 248 |
| MDP-IgG | 2.9 (0.8) | 1.7 – 4.4 | 0.00 (0.00) | - | 0.66 (0.06) | **0.23 (0.11)** | - | 262 |
| MDP-IgM | 3.4 (1.2) | 1.8 – 5.7 | 0.03 (0.15) | - | 0.89 (0.08) | 0.18 (0.14) | - | 268 |
| *Pre-Parturition serum* |  |  |  |  |  |  |  |  |
| KLH-IgA | 2.6 (0.7) | 1.7 – 3.7 | **0.64 (0.16)** | - | 0.36 (0.03) | 0.29 (0.20) | - | 321 |
| KLH-IgG | 4.8 (1.3) | 2.7 – 7.2 | **0.52 (0.16)** | - | 1.38 (0.12) | **0.83 (0.36)** | - | 321 |
| KLH-IgM | 7.0 (1.1) | 5.4 – 8.9 | **0.37 (0.17)** | - | 1.14 (0.10) | 0.15 (0.29) | - | 321 |
| MDP-IgA | 3.3 (1.0) | 1.6 – 4.8 | **0.52 (0.15)** | - | 0.90 (0.08) | **0.48 (0.29)** | - | 320 |
| MDP-IgG | 3.1 (1.1) | 1.5 – 5.0 | 0.25 (0.14) | - | 0.93 (0.08) | **0.44 (0.22)** | - | 321 |
| MDP-IgM | 3.4 (0.9) | 2.0 – 4.9 | **0.40 (0.16)** | - | 0.66 (0.06) | 0.07 (0.22) | - | 321 |

<sup>1</sup>. Significant $h^2$ , $r$ and breed effects are highlighted in bold
<sup>2</sup>. Range shows 5th quantile and 95th quantile, respectively
<sup>3</sup>. Breed effects are relative to Swedish Red (SRB)
<sup>4</sup>. Variance components and effects were estimated using a square root transformation

Heritability estimates for colostrum traits were moderate, ranging from 0.16 to 0.31 and repeatabilities were moderate to high (0.21 to 0.55). Breed effects were significant and consistently higher for SLB and crossbreeds relative to Swedish Red. The effect of breed varied depending on the trait, but the difference between SRB and SLB was about half a standard deviation in most cases. In the case of first test milk, heritability estimates were not significant except for KLH-IgG with a moderate value (0.33). Breed effects, however, were significant for all but one trait, with the difference being approximately one third of a standard deviation. Heritabilities for pre-parturition serum were moderate to high (0.37 to 0.64), except for MDP-IgG which was not significant. SLB had a significantly higher effect on three traits; KLH-IgG, MDP-IgG and KLH-IgM.

Fixed effects in models (1.) and (3.) were selected based on their significance by an incremental Wald F statistics test including interactions. One factor that was tested and not selected was ELISA plate, which was significant by itself, but when the storage plate effect was added it was no longer significant. This was observed both for colostrum and calf serum traits.

For calf serum, tested factors that were not significant include sex, ease of calving, time of first meal and whether the calf was fed with the colostrum of the mother or another cow (as logical variable). For calf serum, volume of first meal and colostrum antibody content was combined as a single “absolute fed colostrum” factor (absolute amount of colostral antibodies fed) with a stronger effect. Figure 1 shows the plots of each calf serum trait versus its matching absolute colostrum trait. Scatter plots show a fairly linear relationship between traits that seems marginally higher for Brix and IgG (0.42 to 0.49) compared to IgM and IgA (0.26 to 0.37).

**Figure 1.**
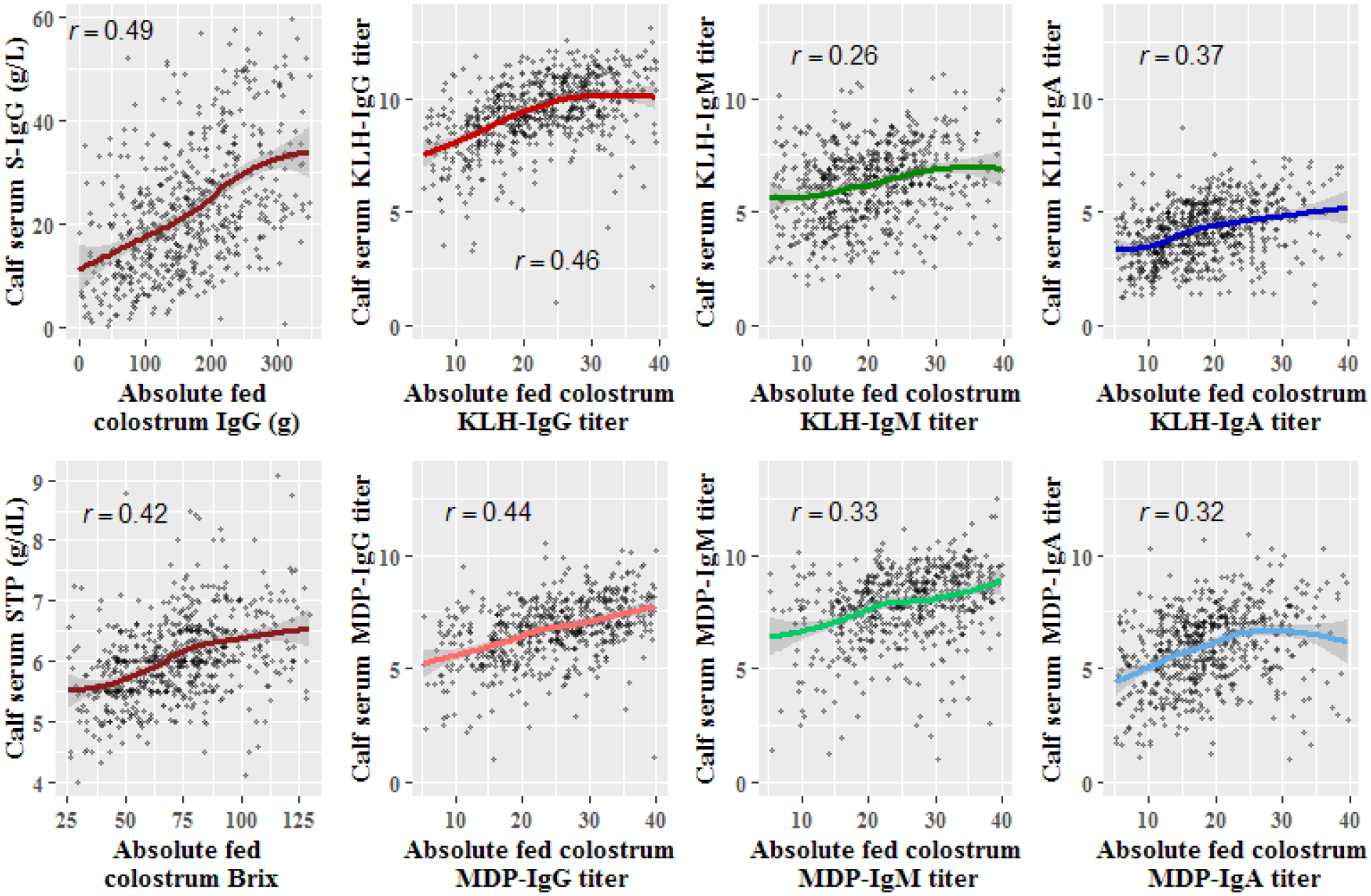
Scatterplots of calf serum traits vs. their matching absolute fed colostrum traits with a LOESS curve and Pearson correlation (r) value.

Time of calving to blood sampling (calves) had varying effects depending on the trait. Figure 2 shows plots for each trait versus the time of blood sampling in days. As opposed to the previous trait, it appears that STP and IgG traits have a slightly lower correlation (−0.09 to −0.21) compared to IgM and IgA (−0.27 to −0.51)

**Figure 2.**
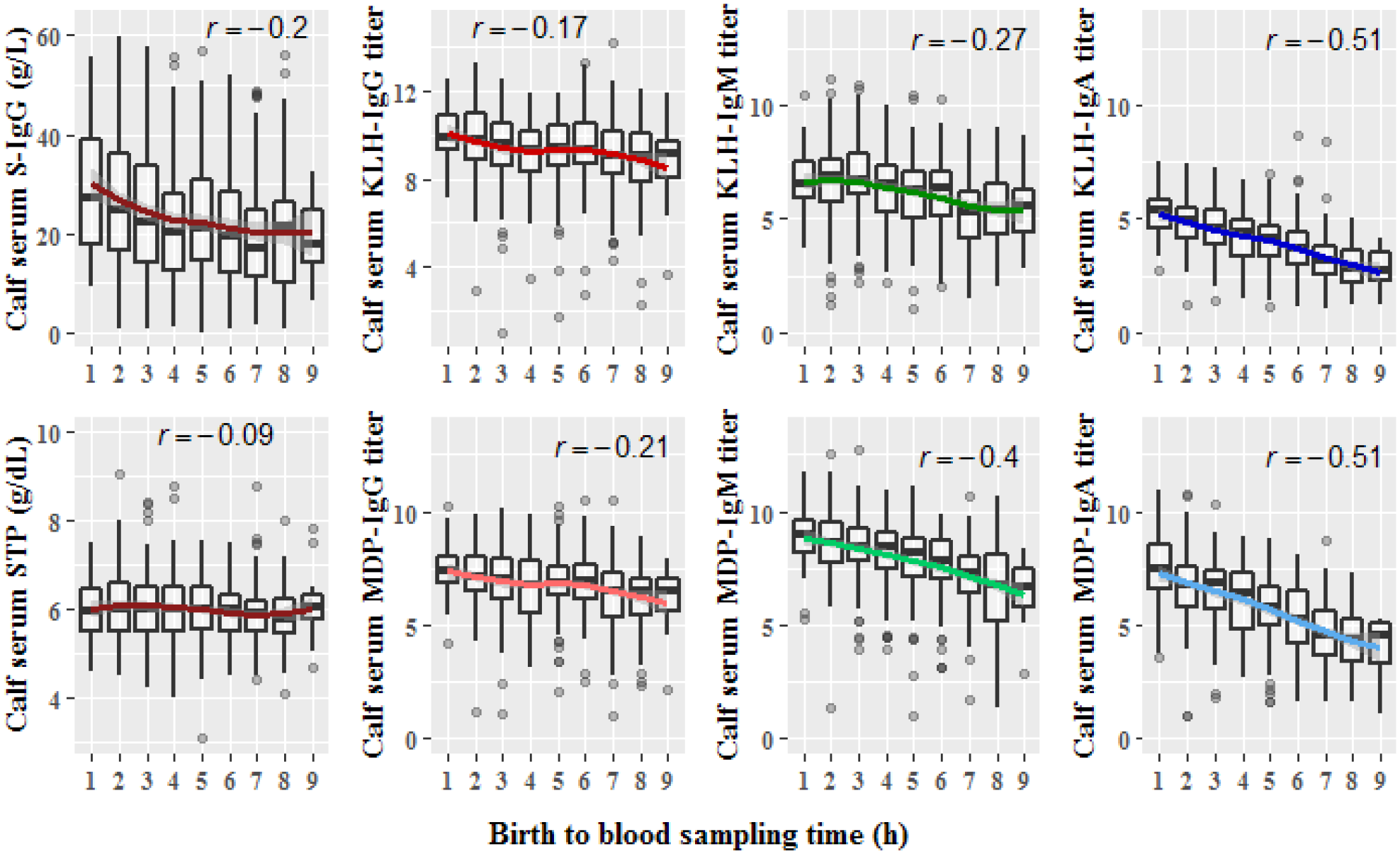
Plots of calf serum traits vs birth to blood sampling time with a LOESS curve and Pearson correlation (r) value.

Genetic correlations for colostrum traits are presented in Table 4. All significant correlations were positive and ranged between 0.49 and 0.97. Most of these were between NAbs of the same isotype or between IgA and IgM NAbs. Brix was positively correlated with all the traits, including total IgG. Similar findings were observed for genetic correlations among calf serum traits (Table 5). Significant correlations ranged from 0.62 to 0.96 and were mostly seen for MDP-IgA and MDP-IgM with other traits, including S-IgG (0.87 and 0.81, respectively).

**Table 4.** Estimated genetic correlations (below diagonal) and phenotypic correlations (above diagonal) between colostrum traits. SE in parenthesis.

| Trait<sup>1</sup> | Brix | T-IgG<sup>2</sup> | KLH-IgA | KLH-IgG | KLH-IgM | MDP-IgA | MDP-IgG | MDP-IgM |
| --- | --- | --- | --- | --- | --- | --- | --- | --- |
| Brix | - | 0.70 (0.02) | 0.52 (0.02) | 0.51 (0.02) | 0.60 (0.02) | 0.48 (0.02) | 0.45 (0.02) | 0.58 (0.02) |
| T-IgG<sup>2</sup> | **0.68 (0.14)** | - | 0.37 (0.04) | 0.51 (0.03) | 0.48 (0.03) | 0.33 (0.04) | 0.35 (0.04) | 0.39 (0.03) |
| KLH-IgA | **0.71 (0.10)** | 0.23 (0.26) | - | 0.55 (0.02) | 0.78 (0.01) | 0.68 (0.01) | 0.45 (0.02) | 0.67 (0.02) |
| KLH-IgG | **0.66 (0.14)** | 0.93 (0.23)<sup>3</sup> | 0.39 (0.21)<sup>3</sup> | - | 0.57 (0.02) | 0.40 (0.02) | 0.56 (0.02) | 0.49 (0.02) |
| KLH-IgM | **0.73 (0.09)** | 0.22 (0.25) | **0.85 (0.06)** | 0.32 (0.23) | - | 0.67 (0.02) | 0.49 (0.02) | 0.79 (0.01) |
| MDP-IgA | **0.52 (0.13)** | 0.05 (0.28) | **0.92 (0.06)** | 0.13 (0.24) | **0.81 (0.09)** | - | 0.41 (0.02) | 0.67 (0.02) |
| MDP-IgG | **0.49 (0.14)** | 0.33 (0.26) | **0.56 (0.15)** | **0.76 (0.15)** | 0.37 (0.18) | 0.44 (0.17)<sup>3</sup> | - | 0.53 (0.02) |
| MDP-IgM | **0.68 (0.10)** | 0.13 (0.27) | **0.82 (0.08)** | 0.10 (0.29) | **0.97 (0.04)** | **0.92 (0.08)** | 0.34 (0.19) | - |

<sup>1</sup>. Significant genetic correlations are highlighted in bold
<sup>2</sup>. Variance components for correlations were estimated using a square root transformation
<sup>3</sup>. Suggestive significance (0.05 < *p*-value < 0.10)

**Table 5.** Estimated genetic correlations (below diagonal) and phenotypic correlations (above diagonal) between calf serum traits. SE in parenthesis.

| Trait<sup>1</sup> | STP | S-IgG<sup>2</sup> | KLH-IgA | KLH-IgG | KLH-IgM | MDP-IgA | MDP-IgG | MDP-IgM |
| --- | --- | --- | --- | --- | --- | --- | --- | --- |
| STP | - | 0.67 (0.02) | 0.49 (0.03) | 0.45 (0.04) | 0.37 (0.04) | 0.48 (0.03) | 0.43 (0.04) | 0.53 (0.03) |
| S-IgG<sup>2</sup> | 0.91 (0.18) | - | 0.57 (0.03) | 0.59 (0.03) | 0.40 (0.04) | 0.60 (0.03) | 0.62 (0.03) | 0.61 (0.03) |
| KLH-IgA | 0.61 (0.25) | 0.70 (0.16)<sup>3</sup> | - | 0.44 (0.04) | 0.53 (0.03) | 0.76 (0.02) | 0.49 (0.03) | 0.71 (0.02) |
| KLH-IgG | 0.71 (0.46) | 0.74 (0.20) | 0.23 (0.25) | - | 0.40 (0.04) | 0.53 (0.03) | 0.66 (0.03) | 0.54 (0.03) |
| KLH-IgM | 0.47 (0.52) | 0.59 (0.28) | 0.80 (0.14)<sup>3</sup> | 0.61 (0.28) | - | 0.51 (0.03) | 0.39 (0.04) | 0.62 (0.03) |
| MDP-IgA | 0.67 (0.31) | **0.87 (0.13)** | **0.91 (0.08)** | 0.86 (0.14)<sup>3</sup> | 0.90 (0.17)<sup>3</sup> | - | 0.59 (0.03) | 0.77 (0.02) |
| MDP-IgG | 0.83 (0.35) | 0.90 (0.15)<sup>3</sup> | 0.80 (0.16)<sup>3</sup> | 0.87 (0.13)<sup>3</sup> | 0.77 (0.25) | 0.99 (0.11) | - | 0.49 (0.03) |
| MDP-IgM | 0.83 (0.26) | **0.81 (0.12)** | **0.84 (0.09)** | **0.62 (0.18)** | **0.91 (0.11)** | **0.96 (0.06)** | 0.77 (0.02)<sup>3</sup> | - |

<sup>1</sup>. Significant genetic correlations are highlighted in bold
<sup>2</sup>. Variance components for correlations were estimated using a square root transformation
<sup>3</sup>. Suggestive significance (0.05 < *p*-value < 0.10)

Correlations were also estimated between colostrum and calf serum traits (Table 6). There were no significant genetic correlations between STP and S-IgG with Brix or colostrum total IgG. Their phenotypic correlations, however, ranged from 0.17 to 0.26. Only IgA and IgM NAbs had significant genetic correlations, which ranged from 0.66 to 0.99.

**Table 6.**
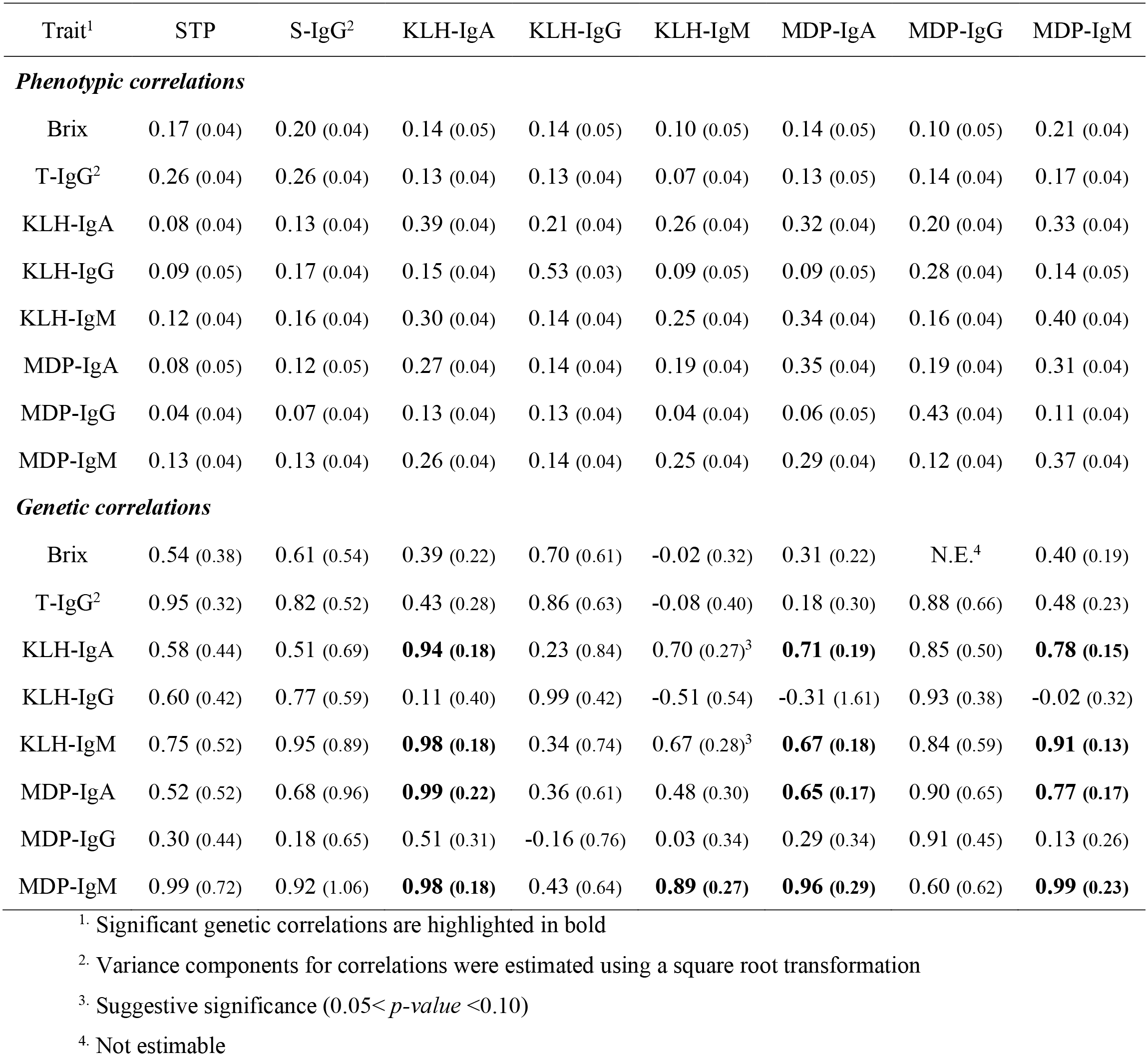
Genetic and phenotypic correlations between colostrum traits (vertical) and calf serum traits (horizontal). SE in parenthesis.

Correlations of colostrum traits with NAbs of first test milk are shown in Table 7. Only colostrum NAbs had significant correlations and ranged from 0.80 to 0.97, mostly between colostrum MDP- IgG and milk NAbs. Genetic correlations were also estimated with pre-parturition serum, having no significant correlations with Brix or total IgG, NAb correlations were between 0.63 and 0.98. Milk production traits for 305d lactation including milk yield (kg), LASCS, fat percentage and protein percentage had a positive trend with colostrum traits, however, none of the traits were significantly correlated.

**Table 7.** Genetic correlations of colostrum traits (horizontal) with first test milk, pre-parturition serum traits and milk production traits. SE in parenthesis.

| Trait<sup>1</sup> | Brix | T-IgG<sup>2</sup> | KLH-IgA | KLH-IgG | KLH-IgM | MDP-IgA | MDP-IgG | MDP-IgM |
| --- | --- | --- | --- | --- | --- | --- | --- | --- |
| *First test milk* |  |  |  |  |  |  |  |  |
| KLH-IgA | 0.14 (0.39) | 0.43 (0.55) | -0.54 (0.51) | 0.93 (0.51) | -0.45 (0.48) | -0.21 (0.41) | **0.86 (0.46)** | -0.07 (0.44) |
| KLH-IgG | 0.66 (0.37) | 0.26 (0.43) | 0.47 (0.36) | 0.79 (0.36) | 0.58 (0.33) | 0.41 (0.35) | **0.98 (0.37)** | 0.43 (0.34) |
| KLH-IgM | 0.83 (0.84) | 0.68 (0.89) | 0.99 (0.75) | 0.78 (0.40) | **0.87 (0.20)** | **0.84 (0.24)** | **0.80 (0.30)** | **0.97 (0.21)** |
| MDP-IgA | 0.02 (0.37) | -0.22 (0.53) | 0.59 (0.35) | 0.82 (0.71) | 0.50 (0.43) | 0.59 (0.56) | 1.00 (0.70) | 0.47 (0.45) |
| MDP-IgG | 0.86 (0.64) | 0.61 (1.25) | 0.83 (0.49) | 0.83 (0.64) | 0.75 (0.28)<sup>3</sup> | 0.59 (0.33) | **0.97 (0.30)** | 0.59 (0.67) |
| MDP-IgM | -0.25 (4.68) | -0.24 (4.79) | 1.00 (0.99) | N.E.<sup>4</sup> | **0.98 (0.29)** | 0.72 (0.44) | N.E.<sup>4</sup> | 0.92 (1.40) |
| *Pre-parturition serum* |  |  |  |  |  |  |  |  |
| KLH-IgA | -0.21 (0.23) | -0.13 (0.30) | 0.40 (0.23) | 0.20 (0.32) | **0.63 (0.23)** | **0.92 (0.17)** | 0.55 (0.26)<sup>3</sup> | **0.80 (0.19)** |
| KLH-IgG | -0.12 (0.23) | 0.30 (0.31) | -0.02 (0.25) | 0.60 (0.36) | -0.40 (0.29) | -0.22 (0.26) | 0.43 (0.26) | -0.44 (0.26) |
| KLH-IgM | -0.19 (0.32) | -0.59 (0.42) | 0.48 (0.36) | 0.45 (0.49) | 0.58 (0.38) | 0.94 (0.52)<sup>3</sup> | 0.45 (0.39) | **0.85 (0.33)** |
| MDP-IgA | -0.28 (0.23) | 0.03 (0.31) | 0.45 (0.24) | 0.23 (0.33) | 0.39 (0.25) | **0.79 (0.21)** | **0.98 (0.24)** | **0.68 (0.24)** |
| MDP-IgG | 0.04 (0.33) | 0.35 (0.43) | **0.70 (0.32)** | **0.99 (0.35)** | -0.03 (0.35) | 0.41 (0.31) | **0.82 (0.35)** | -0.16 (0.37) |
| MDP-IgM | -0.43 (0.29) | -0.30 (0.35) | 0.43 (0.30) | -0.04 (0.44) | 0.36 (0.31) | **0.89 (0.30)** | 0.23 (0.33) | **0.79 (0.31)** |
| *Milk production* |  |  |  |  |  |  |  |  |
| Milk yield | 0.25 (0.17) | -0.02 (0.28) | 0.04 (0.18) | 0.36 (0.26) | -0.07 (0.19) | 0.12 (0.18) | 0.15 (0.20) | -0.10 (0.20) |
| LASCS | N.E.<sup>4</sup> | 0.03 (0.29) | N.E.<sup>4</sup> | N.E.<sup>4</sup> | N.E.<sup>4</sup> | -0.42 (0.87) | 0.21 (0.27) | -0.15 (0.24) |
| Fat % | -0.15 (0.13) | 0.15 (0.21) | N.E.<sup>4</sup> | N.E.<sup>4</sup> | N.E.<sup>4</sup> | 0.00 (0.14) | 0.09 (0.15) | 0.08 (0.15) |
| Protein % | 0.00 (0.12) | 0.23 (0.20) | N.E.<sup>4</sup> | N.E.<sup>4</sup> | N.E.<sup>4</sup> | 0.08 (0.13) | 0.14 (0.14) | 0.13 (0.14) |

<sup>1</sup>. Significant genetic correlations are highlighted in bold
<sup>2</sup>. Variance components for correlations were estimated using a square root transformation
<sup>3</sup>. Suggestive significance (0.05 < *p*-value < 0.10)
<sup>4</sup>. Not estimable

Correlations of calf serum antibody levels with health and production traits are presented in Table 8. Only LASCS had significant correlations with calf serum antibody levels, ranging from −0.66 to −0.98 for IgM and MDP-IgA. Calf health had no significant genetic correlations, but it had a borderline significant (*p* = 0.08) negative genetic correlation with MDP-IgG and a consistent negative trend can be seen in calf health versus all the other antibody traits, both regarding phenotypic and genetic correlations.

**Table 8.** Genetic and phenotypic correlations of calf serum traits with milk production and health traits.

| Trait<sup>1, 2</sup> | STP | S-IgG<sup>3</sup> | KLH-IgA | KLH-IgG | KLH-IgM | MDP-IgA | MDP-IgG | MDP-IgM |
| --- | --- | --- | --- | --- | --- | --- | --- | --- |
| *Phenotypic correlations* |  |  |  |  |  |  |  |  |
| Calf health | -0.04 (0.06) | -0.11 (0.07) | -0.08 (0.06) | -0.05 (0.06) | -0.12 (0.07) | -0.12 (0.06) | -0.12 (0.06) | -0.12 (0.07) |
| Adult health | -0.04 (0.05) | -0.06 (0.05) | 0.00 (0.05) | -0.04 (0.05) | 0.04 (0.05) | -0.08 (0.05) | -0.02 (0.05) | 0.03 (0.05) |
| AFC | -0.04 (0.08) | 0.03 (0.08) | -0.13 (0.08) | -0.16 (0.08) | 0.03 (0.08) | 0.05 (0.08) | -0.01 (0.08) | -0.11 (0.08) |
| Fat % | -0.12 (0.07) | -0.02 (0.07) | -0.02 (0.07) | -0.09 (0.07) | -0.03 (0.07) | -0.08 (0.07) | -0.04 (0.07) | -0.05 (0.07) |
| Milk yield | -0.03 (0.07) | -0.06 (0.08) | -0.04 (0.07) | -0.07 (0.07) | -0.09 (0.08) | -0.11 (0.07) | -0.05 (0.08) | -0.11 (0.08) |
| Protein % | 0.02 (0.07) | 0.06 (0.07) | 0.02 (0.07) | -0.01 (0.07) | 0.06 (0.07) | 0.05 (0.07) | 0.05 (0.07) | 0.02 (0.07) |
| LASCS | -0.09 (0.07) | -0.12 (0.07) | 0.01 (0.07) | 0.05 (0.07) | -0.12 (0.07) | -0.19 (0.07) | -0.10 (0.07) | 0.07 (0.07) |
| *Genetic correlations* |  |  |  |  |  |  |  |  |
| Calf health | 0.01 (0.75) | -0.42 (0.58) | -0.52 (0.47) | -0.72 (0.81) | -0.74 (0.58) | -0.53 (0.41) | -0.93 (0.41)<sup>4</sup> | -0.77 (0.47) |
| Adult health | 1.00 (1.43) | 0.36 (0.63) | 0.87 (1.05) | 0.14 (0.61) | 0.74 (0.76) | 0.44 (0.59) | 0.68 (0.89) | 0.08 (0.35) |
| AFC | -0.18 (0.25) | 0.09 (0.18) | -0.27 (0.16) | -0.44 (0.18)<sup>4</sup> | -0.06 (0.19) | -0.02 (0.14) | -0.09 (0.16) | -0.23 (0.16) |
| Fat % | -0.18 (0.22) | 0.00 (0.18) | -0.14 (0.16) | -0.08 (0.17) | 0.06 (0.19) | -0.13 (0.14) | -0.16 (0.17) | -0.11 (0.15) |
| Milk yield | -0.38 (0.27) | -0.20 (0.20) | 0.00 (0.18) | 0.06 (0.20) | -0.20 (0.21) | -0.32 (0.16) | -0.16 (0.19) | -0.28 (0.17) |
| Protein % | 0.21 (0.21) | 0.21 (0.17) | -0.06 (0.16) | -0.16 (0.18) | 0.06 (0.18) | 0.10 (0.14) | 0.18 (0.16) | 0.12 (0.15) |
| LASCS | -0.52 (0.33) | -0.62 (0.26)<sup>4</sup> | -0.38 (0.25) | -0.63 (0.26)<sup>4</sup> | **-0.98 (0.26)** | **-0.66 (0.22)** | -0.74 (0.30)<sup>4</sup> | **-0.86 (0.23)** |

<sup>1.</sup> Significant genetic correlations are highlighted in bold837 <sup>2.</sup> Calf health refers to the Healthy/Disease data for calves between 0 to 3 months of age, Adult health refers to the Healthy/Disease data for animals between 8 to 36 months of age839 <sup>3.</sup> Variance components for correlations were estimated using a square root transformation840 <sup>4.</sup> Suggestive significance (0.05 < *p-value* < 0.10)

## Discussion

In this study we found that there is a significant genetic contribution to variation for most of the traits in both colostrum and calf serum, as well as genetic correlations between these traits. In addition, variation in most of the calf serum traits was substantially affected by environmental differences between mothers.

### Heritabilities

Two indicator traits were analyzed; Brix for colostrum and STP for calf serum. They both approximate antibody content by quantifying total solids (TS) and provide methodologically simpler but less accurate measurements compared to ELISA or radial immunodiffusion (**RID**). Brix had a moderate heritability (0.31), which was similar to the value reported (0.27) in a previous colostrum study [8]. In contrast, heritability for STP was not significant, which could be due to the variation from the rest of the molecules that are co-measured masking the genetic variance of the antibody level.

Heritabilities for IgG concentration (g/L) were 0.20 and 0.25 for colostrum and calf serum, respectively. In both cases, a square root transformation was applied to normalize the residual distribution. A previous study reported a heritability of 0.10 for colostrum IgG in Irish Holstein and crossbreeds [28]. Although the difference between the studies can be explained by different populations and breeds, it suggests that breeding for higher colostrum IgG levels is indeed possible.

In the case of calf serum, our results concur with the findings of Gilbert et al. (1988) and Martin et al. (2021) who estimated significant heritabilibites for S-IgG, confirming that it is possible to also breed for increased antibody uptake from colostrum by calves.

Two antigens were used to measure NAbs: KLH and MDP. The latter is a microbial pathogen-associated molecular pattern (**PAMP**) comprising the minimal peptidoglycan (**PGN**) motif common to both Gram-positive and Gram-negative bacteria [29] such as *Escherichia coli* and *Staphylococcus aureus*, which are ubiquitous in most environments. In previous studies, antibodies to PGN have been regarded as NAbs reflecting the animals’ innate humoral response [30, 31]. Given the universal presence of MDP (PGN) it is, however, reasonable to assume that the measurements for this antigen could partly represent antibodies produced by the cows as part of adaptive immune response to bacteria in the environment. KLH, on the other hand, is considered as a true measure of NAbs, since cows are not normally exposed to this antigen [32]. For colostrum, all NAb heritabilities were significant and for five out the six traits ranged from 0.22 to 0.29, with the exception of KLH-IgG (0.16). This range is within the values reported before for cow serum and milk [30, 33]. There were no major differences between isotype heritabilities except KLH-IgG. In the case of calf serum NAbs, heritabilities ranged from 0.23 to 0.59. NAbs of IgA isotype had almost identical heritabilities (0.43 and 0.46) which was observed for IgG (0.24 and 0.26) as well. However, there was a big difference in heritability between KLH-IgM (0.23) and MDP-IgM (0.59). This difference in IgM NAbs suggests that the way they are absorbed may be affected by the antigens they bind (or not bind to) and since IgM molecules are much larger than IgG and IgA molecules, different transportation mechanisms may be involved.

For first test milk, NAb heritability estimates were not significant for most traits, most likely due to the limited number of samples (standard errors ranged from 0.15 to 0.20). Other studies have estimated heritabilities for KLH NAbs of different isotypes in milk [30, 33] with values ranging from 0.08 to 0.48. In the case of pre-parturition serum, in spite of the large standard errors, heritabilities were significant except for MDP-IgG. There are no reported heritabilities for MDP-IgG in bovine serum, but a previous study in milk [30] estimated a value of 0.15 for PGN-IgG1.

Our results show moderate heritabilities for colostrum traits, including Brix (0.31) as an indicator, implying that genetic selection for colostrum quality is possible. Brix has the advantage of being easier to perform than an ELISA or RID and can potentially be implemented on farms for routine assessment of colostrum quality. For calf traits, heritability estimates were moderate to high, showing that there is an important genetic component for these traits, and that the occurrence of FPT can potentially be reduced through genetic selection. Unfortunately, the heritability for STP, the indicator trait for calf serum, was not significant. Providing a practical way to measure absorption of colostrum antibodies by the calf is critical to effectively implement this trait in a breeding program. For future studies, it could be interesting to analyze calf serum using a Brix refractometer since some studies show a high correlation with serum IgG [34].

### Repeatabilities (colostrum)

NAb in colostrum had notably higher permanent environment effects (0.10 to 0.31) than Brix (0.04) and total IgG (0.01) which made their repeatabilities proportionally higher. Permanent environment effect in this case was the effect of repeated measures from the same cow in different parities: the higher the measurements correlate across parities, the stronger the effect. The observed differences are probably due to an accumulated exposure to antigens in older cows, which generates more specific antibodies that get transferred from the serum to colostrum [28]. NAb, on the other hand, are maintained largely constant throughout life [35].

### Maternal contribution (calf serum)

The maternal contribution to variation in calf serum traits ranged from 17 to 37% of the variance, except for KLH-IgA, which showed no significant maternal effect. Maternal effect in this case was the non-genetic contribution of the dam across different calvings on calf serum traits, the more similar the measurements between maternal siblings, the stronger the effect. For calf serum antibodies, we believe that the biggest (non-genetic) maternal contribution may come from the colostrum, and since we accounted for colostrum antibodies in the model, other colostrum components could be influencing how well the calf absorbs antibodies. Some components of colostrum are fat, protein, peptides, non-protein nitrogen, vitamins and minerals, hormones, growth factors, cytokines and nucleotides [36]. It seems plausible that some of these components differ between colostrum of different cows and directly or indirectly affect antibody uptake. One example is lactoferrin, an iron-sequestering glycoprotein with antimicrobial, anti-inflammatory, and anti-oxidative properties [37]. By inhibiting bacterial growth it might indirectly influence IgG uptake.

Aside from colostrum, there is increasing evidence that the calf intestine is not sterile until birth. A study by Alipour et al. (2018) found a low-abundant microbiota in rectal meconium and mucosa of calves sampled at birth resembling the composition of dam oral microbiota, but included typical intestinal taxa. The exact mechanism of how these bacteria colonize *in utero* is not clear, but these results suggest another source of maternal effect that could impact antibody uptake in the calf.

### Breed

Our results show that the effect of breed is slightly higher for SLB compared to SRB or CRB in colostrum, first test milk and pre-parturition serum traits. Different sample types were analyzed by different techniques, but in all cases SLB had higher values. However, even for statistically significant effects, most *p-values* were barely below 0.05. Holsteins are generally assumed to have a poorer colostrum quality, but several studies have found non-significant differences between Holstein and other breeds concerning colostrum IgG [39–41]. Breed effects were not significant for calf serum traits except STP, for which the same pattern was seen as in colostrum.

### Other effects

Time of first meal is a critical factor to avoid FPT [42], but in our case it was not significant. This is probably due to the fact that feedings were done within the appropriate window of time, as 95% of the first meals were given before 6 hours after birth (not shown) which is the cut-off point for optimal feeding time [43] and significantly below the 24h cut-off.

Sampling time after birth showed a stronger negative correlation for IgM and IgA traits compared to IgG traits and STP. This is most likely because IgG has a half-life of 28.5 days in colostrum-fed calves [44], as opposed to IgA and IgM with only 2.8 and 4.8 days respectively [45], causing a more pronounced decline in time for these two. This could also explain what was seen for calf serum concentration versus absolute fed colostrum (Figure 1.), where the trend was reversed. IgG traits and STP had a positive slope (0.45) and IgA and IgM a less pronounced one (0.32) because of their shorter half-lives.

### Genetic correlations

#### Colostrum traits

Brix was significantly and positively correlated with all the colostrum antibody traits, ranging from 0.49 to 0.73. Several studies have pointed out the use of Brix to approximate the amount of antibodies in colostrum [6, 46], namely IgG, but to the best of our knowledge this is the first report of genetic correlation with total IgG. Specifically, the correlation between Brix and total IgG was 0.68, and even if the response to selection for Brix may be lower, it should be possible to collect more observations for this trait given its technical practicality and possibility to measure at the farm compared to an ELISA or RID test (which measure total IgG directly), making it a promising indicator trait for selection of higher quality colostrum.

Aside from Brix, total IgG was not genetically correlated with any other trait, but with KLH-IgG the correlation of 0.93 (0.23) was close to being significantly different from 0 (*p* = 0.06). NAb traits showed a correlation pattern similar to that observed in a previous study of NAbs in milk [30], where IgM and IgA traits had very strong and positive genetic correlations between and within isotypes (0.82 to 0.97), suggesting a common genetic background for IgA and IgM. KLH-IgG and MDP-IgG had a correlation of 0.76, which is high but low enough to assert that they are different traits.

#### Calf serum traits

Unlike colostrum, the indicator trait for calf serum (STP), did not have significant genetic correlations with any trait. This is due to the non-significant genetic variance and covariance of this trait. S-IgG had significant genetic correlations only with MDP-IgA and MDP-IgM and a suggestive significant correlation (*p* = 0.08) with MDP-IgG. Interestingly, MDP-IgM had positive genetic correlations with all of the traits except STP and MDP-IgG, the latter having a suggestive significant association. MDP- IgA also had significant or suggestive significant genetic correlations with most traits except STP and MDP-IgG. Even though there is a slightly higher genetic correlation between traits from the same isotype and between IgA and IgM in calf serum, it is not as pronounced as in colostrum. This implies that isotype may play a less important role for uptake which is something that Burton et al. (1989) observed as well.

#### Colostrum traits versus calf serum traits

Measuring antibody absorption by the calf requires taking blood samples from the animal and centrifuging them to separate the serum. This means that unlike colostrum that can be sampled and measured on the farm (using a Brix refractometer), calf serum analyses require a veterinarian or technician for the sampling and a laboratory setting. For this reason, we wanted to estimate if colostrum quality in the cow correlates genetically with the calf’s ability to absorb antibodies.

Originally, we attempted to correlate colostrum traits using cow for the genetic component with calf serum traits using calf for the genetic component, but there were convergence problems and correlations could not be estimated. Instead, we estimated these correlations using cow for the genetic effect on colostrum and calf serum traits. This is not ideal, but we wanted to get an idea of how these traits might correlate.

Only NAb IgA and IgM traits had significant genetic correlations within and between isotype traits. It seems that only traits with very strong correlations could pass the significance threshold and even then the values had large standard errors (0.13 to 0.29). In spite of most correlations not being significant, a positive tendency could be seen for some of them, such as colostrum Brix and total IgG with calf serum IgG.

A larger sample size is necessary to estimate these correlations more accurately, and to properly correlate the colostrum traits (using cow for the genetic effect) with the calf serum traits using calf for the genetic component, instead of cow like in this study.

#### Colostrum traits versus first test milk and pre-parturition serum

Neither colostrum Brix nor total IgG were significantly correlated with any of the first test milk or pre-parturition serum NAb traits. A pattern was not apparent from the significant genetic correlations for both first test milk and pre-parturition serum NAbs, which may be due to the small number of milk and serum samples used for the analysis (<300).

### Production traits

#### Colostrum

To test if genetic selection for colostrum antibodies may have a negative effect on milk production, we estimated genetic correlations with milk yield, fat and protein percentage, and lactation average somatic cell score for 305 d lactation period following the calving at which the colostrum was sampled. We found no significant genetic correlations between colostrum traits and milk production, although standard errors were rather high (0.13 to 0.28). Nonetheless, results suggest that there are no strong genetic correlations. Total IgG tended to be positively correlated with fat and protein percentage. Brix and protein yield seemed to have a positive trend, whereas Brix and fat percentage had a slightly negative trend. Genetic correlation could not be estimated for Brix and LASCS, but phenotypic correlation was 0.12 (0.05).

#### Calves

There is an unclear relationship between calf serum IgG (or FPT) and milk production performance later in life. Two studies usually referenced for this topic are: a) Faber et al. (2005) who found that calves fed 2L of high quality colostrum at birth produced significantly less milk during the first and second lactations compared to animals that were fed 4L and b) DeNise et al. (1989) who reported that for every additional g/L of IgG in serum of calves from 24 to 48 hours of age, an increase of 8.5 kg of milk and 0.24 kg of fat was observed during the first lactation. However, we were not able to find additional studies that replicate these results or even follow-ups from the same groups. Our analysis did not find significant genetic correlations between calf serum traits and milk yield, protein percentage or fat percentage.

Lactation average SCS is a log-derived measurement of SCC, a trait that is used as a surrogate for clinical mastitis and overall udder health. LASCS has a good genetic correlation with clinical mastitis (0.6 to 0.7) [49, 50]. In our study, we found significant negative genetic correlations between NAbs and LASCS. KLH-IgM, MDP-IgM and MDP-IgA had correlations of −0.98, −0.96 and −0.66 respectively. Additionally, MDP-IgG, KLH-IgG and S-IgG had borderline significant negative correlations. These findings suggest that NAbs could be used to select for animals less prone to clinical mastitis, agreeing with the results found by Thompson-Crispi et al. (2013) for KLH-IgM and Ploegaert et al. (2010) for IgM and IgA NAbs. Further analyses with a larger dataset are needed to confirm this association.

We did not find significant genetic correlations of calf serum traits with age at first calving (AFC). KLH-IgG, however, had a borderline significant negative correlation with AFC (–0.44), which is the desired direction, since increasing KLH-IgG would lead to lower AFC. Age at first calving has been described as a proxy for average daily gain (**ADG**) since a higher value leads to an earlier insemination [2]. This result is in agreement with Furman-Fratczak et al. (2011) who found that calves with higher IgG in serum had higher growth rates allowing for earlier inseminations.

### Calf health traits

Health information from Röbäcksdalen calves from 0 to 3 months of age should reflect a period where maternal passive immunity still influences the health of the calves. For this trait, we had 63 reported cases out of 231 animals, primarily pneumonia (70%) and diarrhea (15%), where 61 of the cases were treated with antibiotics, implying bacterial infection. This means that the binary 0/1 trait is a proxy for healthy/bacterial infection. Genetic correlations for this trait were not significant, but a negative tendency was observed for all serum traits except STP. This is a favorable trend, since the higher the antibody titer, the value will lean more towards healthy. MDP-IgG had a suggestive significant negative genetic correlation (−0.93) but with a rather large standard error (0.41). MDP is known to stimulate the pattern recognition receptor known as Nucleotide-binding oligomerization domain-containing protein 2 (**NOD2**). This receptor then activates the protein complex NF-κB (nuclear factor kappa-light-chain-enhancer of activated B cells) [29]. Hence, it is possible that animals with higher levels of MDP-IgG are able to prime better the immune system by recognizing bacterial PAMPs and triggering NF-κB [53].

Data for animals from 8 to 36 months should reflect long term health effects of maternal antibody levels in the calves’ serum. We had 66 cases among 301 animals from two farms for this trait. Most cases had an unknown etiology (47%), followed by infectious (27%) and metabolic (22%) diseases. Given the small number of cases with known etiologies, it was not possible to subcategorize the trait and it was coded as 0/1 for healthy/sick. No genetic correlations were observed for any of the traits, and most trends observed were obscured by large standard errors.

Given the large standard errors observed for health traits, it might be possible to estimate more accurately these correlations with a larger number of samples.

## Conclusions

We have shown that all but one of the measured calf serum traits are heritable, including S-IgG, indicating that genetic selection can be used to reduce FPT. Unfortunately, STP was not heritable and an indicator trait for calf serum that is easy to measure is key to use this trait in breeding programs. Interestingly, we found that there is a significant maternal contribution to calf serum antibody content beyond colostrum antibodies. Also, further analyses are needed to establish the genetic relationship between calf serum antibodies and health later in life.

Brix in colostrum has positive genetic correlations with the antibody traits investigated, thus Brix can be used as an indicator trait for selection of higher quality colostrum. Also, Brix is not genetically correlated with milk production traits in an unfavorable way.

Our results suggest that these traits can be used for selection programs focused on genetically improving antibody content in both colostrum and calf serum, pending a practical indicator trait for calf S-IgG.

## List of abbreviations

ADG: Average daily gain
AFC: Age at first calving
CRB: Swedish Red and Swedish Holstein crossbreds
EDTA: Ethylenediaminetetraacetic acid
ELISA: Enzyme-linked immunosorbent assay
FPT: Failure of passive transfer
HRP: Horseradish peroxidase
Ig: Immunoglobulin
KLH: Keyhole limpet hemocyanin
LASCS: Lactation-average Somatic Cell Score
lpw: Last positive well
MDP: Muramyl dipeptide
NAb: Natural antibodies
NF-κB: Nuclear factor kappa-light-chain-enhancer of activated B cells
NOD2: Nucleotide-binding oligomerization domain-containing protein 2
OD: Optical density
PAMP: Pathogen-associated molecular pattern (PAMP)
PBST: Phosphate-buffered saline with Tween 20
PGN: Peptidoglycan
RID: Radial immunodiffusion
SCC: Somatic Cell Count
SLB: Swedish Holstein
SRB: Swedish Red
STP: Serum total protein
S-IgG: Serum IgG
TMC: Tetramethylbenzidine
TS: Total solids

## Declarations

### Ethics approval and consent to participate

All animal care and handling procedures used in this study were reviewed and approved by the Swedish Ethical Committee on Animal Research (Uppsala djurförsöksetiska nämnd), approval number C 140/14. The animals in Röbäcksdalen were handled and kept with permission from the Swedish Ethical Committee on Animal Research represented by the Court of Appeal for Northern Norrland in Umeå, Sweden by the approval A 17/2016.

### Consent for publication

Not applicable.

### Availability of data and materials

Some of the data that support the findings of this study were provided by Växa Sverige but restrictions apply to the availability of these data, which were used under license for the current study, and so are not publicly available. Data are however available from the authors upon reasonable request and with permission of Växa Sverige.

### Authors’ contributions

JCS performed the research, analysis, data interpretation and wrote the manuscript. TdH, MJ, AL and AM performed the research, analysis and data interpretation. JJW, MT, DJK and HB designed the experiment and overall study, interpreted the data and critically revised the manuscript. JA and HKP analyzed and interpreted the data and revised the manuscript. In addition to their aforemention contributions, JJW and JA also performed the research. All authors read and approved the final manuscript.

## Acknowledgements

We would like to thank the staff at the experimental farms of Lövsta, Röbäcksdalen SITES (Dr Mårten Hetta, Annika Moström, Gun Bernes and Reija Danielsson) and Viken. Special thanks to Hans Stålhammar from Viking genetics for all the help with the animal IDs and Ger de Vries Reilingh for the assistance in analyzing the colostrum samples. The part of the study that has been conducted in Röbäcksdalen, Umeå has been made possible by the use of the Swedish Infrastructure for Ecosystem Science (SITES), in this case by support of the field station Röbäcksdalen.

## References

1. Weaver DM, Tyler JW, VanMetre DC, Hostetler DE, Barrington GM (2000) Passive transfer of colostral immunoglobulins in calves. J Vet Intern Med 14:569–77

2. Raboisson D, Trillat P, Cahuzac C (2016) Failure of Passive Immune Transfer in Calves: A Meta-Analysis on the Consequences and Assessment of the Economic Impact. PLoS One 11:e0150452

3. Liberg P (2000) Råmjölksutfodring - “En god start förlänger livet” (Colostrum feeding - “A good start prolongs life”). Proceedings of the Svenskt Veterinärrnöte, 9-10 November, Uppsala, Sweden, pp 133–139 (In Swedish)

4. Torsein M, Lindberg A, Sandgren CH, Waller KP, Törnquist M, Svensson C (2011) Risk factors for calf mortality in large Swedish dairy herds. Prev Vet Med 99:136–147

5. Bielmann V, Gillan J, Perkins NR, Skidmore AL, Godden S, Leslie KE (2010) An evaluation of Brix refractometry instruments for measurement of colostrum quality in dairy cattle. J Dairy Sci 93:3713–3721

6. Løkke MM, Engelbrecht R, Wiking L (2016) Covariance structures of fat and protein influence the estimation of IgG in bovine colostrum. J Dairy Res 83:58–66

7. Gilbert RP, Gaskins CT, Hillers JK, Brinks JS, Denham AH (1988) Inbreeding and immunoglobulin G1 concentrations in cattle. J Anim Sci 66:2490–7

8. Soufleri A, Banos G, Panousis N, Fletouris D, Arsenos G, Valergakis GE (2019) Genetic parameters of colostrum traits in Holstein dairy cows. J Dairy Sci 102:11225–11232

9. Nilsson D (2015) Factors of importance for high vs low uptake of immunoglobuline from colostrum in calves. Swedish University of Agricultural Sciences. http://urn.kb.se/resolve?urn=urn:nbn:se:slu:epsilon-s-4619

10. Burton JL, Kennedy BW, Burnside EB, Wilkie BN, Burton JH (1989) Variation in serum concentrations of immunoglobulins G, A, and M in Canadian Holstein-Friesian calves. J Dairy Sci 72:135–49

11. Martin P, Vinet A, Denis C, Grohs C, Chanteloup L, Dozias D, Maupetit D, Sapa J, Renand G, Blanc F (2021) Determination of immunoglobulin concentrations and genetic parameters for colostrum and calf serum in Charolais animals. J Dairy Sci 104:3240–3249

12. Svensson C, Linder A, Olsson S-O (2006) Mortality in Swedish Dairy Calves and Replacement Heifers. J Dairy Sci 89:4769–4777

13. Autio T, Pohjanvirta T, Holopainen R, Rikula U, Pentikäinen J, Huovilainen A, Rusanen H, Soveri T, Sihvonen L, Pelkonen S (2007) Etiology of respiratory disease in non-vaccinated, non-medicated calves in rearing herds. Vet Microbiol 119:256–265

14. Closs G, Dechow C (2017) The effect of calf-hood pneumonia on heifer survival and subsequent performance. Livest Sci 205:5–9

15. Avrameas S (1991) Natural autoantibodies: from “horror autotoxicus” to “gnothi seauton”. Immunol Today 12:154–9

16. Reyneveld GIj, Savelkoul HFJ, Parmentier HK (2020) Current Understanding of Natural Antibodies and Exploring the Possibilities of Modulation Using Veterinary Models. A Review. Front Immunol 11:2139

17. Thompson-Crispi KA, Miglior F, Mallard BA (2013) Genetic parameters for natural antibodies and associations with specific antibody and mastitis in Canadian Holsteins. J Dairy Sci 96:3965–72

18. de Klerk B (2016) Antibodies and Longevity of Dairy Cattle: Genetic Analysis. PhD Thesis. Wageningen University

19. Denholm SJ, McNeilly TN, Banos G, Coffey MP, Russell GC, Bagnall A, Mitchell MC, Wall E (2018) Immune-associated traits measured in milk of Holstein-Friesian cows as proxies for blood serum measurements. J Dairy Sci 101:10248–10258

20. Machado VS, Bicalho MLS, Gilbert RO, Bicalho RC (2014) Short communication: Relationship between natural antibodies and postpartum uterine health in dairy cows. J Dairy Sci 97:7674–8

21. Star L, Frankena K, Kemp B, Nieuwland MGB, Parmentier HK (2007) Natural humoral immune competence and survival in layers. Poult Sci 86:1090–9

22. Berghof TVL, Matthijs MGR, Arts JAJ, Bovenhuis H, Dwars RM, van der Poel JJ, Visker MHPW, Parmentier HK (2019) Selective breeding for high natural antibody level increases resistance to avian pathogenic Escherichia coli (APEC) in chickens. Dev Comp Immunol 93:45–57

23. Wijga S, Bastiaansen JWM, Wall E, Strandberg E, de Haas Y, Giblin L, Bovenhuis H (2012) Genomic associations with somatic cell score in first-lactation Holstein cows. J Dairy Sci 95:899–908

24. Ploegaert TCW, Wijga S, Tijhaar E, van der Poel JJ, Lam TJGM, Savelkoul HFJ, Parmentier HK, van Arendonk J a. M (2010) Genetic variation of natural antibodies in milk of Dutch Holstein-Friesian cows. J Dairy Sci 93:5467–5473

25. Frankena K (1987) The interaction between Cooperia spp. and Ostertagia spp. (Nematoda: Trichostrongylidae) in cattle. Wageningen University

26. de Koning DB, Damen EPCW, Nieuwland MGB, van Grevenhof EM, Hazeleger W, Kemp B, Parmentier HK (2015) Association of natural (auto-) antibodies in young gilts with osteochondrosis at slaughter. Livest Sci 176:152–160

27. Gilmour AR, Gogel BJ, Cullis BR, Welham SJ, Thompson R (2015) ASReml user guide release 4.1. VSN Int Ltd 364

28. Conneely M, Berry DP, Sayers R, Murphy JP, Lorenz I, Doherty ML, Kennedy E (2013) Factors associated with the concentration of immunoglobulin G in the colostrum of dairy cows. Animal 7:1824–32

29. Girardin SE, Boneca IG, Viala J, Chamaillard M, Labigne A, Thomas G, Philpott DJ, Sansonetti PJ (2003) Nod2 is a general sensor of peptidoglycan through muramyl dipeptide (MDP) detection. J Biol Chem 278:8869–8872

30. Wijga S, Bovenhuis H, Bastiaansen JWM, van Arendonk J a. M, Ploegaert TCW, Tijhaar E, van der Poel JJ (2013) Genetic parameters for natural antibody isotype titers in milk of Dutch Holstein-Friesians. Anim Genet 44:485–492

31. Ploegaert TCW, Tijhaar E, Lam TJGM, Taverne-Thiele A, van der Poel JJ, van Arendonk J a M, Savelkoul HFJ, Parmentier HK (2011) Natural antibodies in bovine milk and blood plasma: variability among cows, repeatability within cows, and relation between milk and plasma titers. Vet Immunol Immunopathol 144:88–94

32. van Knegsel ATM, de Vries Reilingh G, Meulenberg S, van den Brand H, Dijkstra J, Kemp B, Parmentier HK (2007) Natural Antibodies Related to Energy Balance in Early Lactation Dairy Cows. J Dairy Sci 90:5490–5498

33. de Klerk B, Ducro BJ, Heuven HCM, den Uyl I, van Arendonk JAM, Parmentier HK, van der Poel JJ (2015) Phenotypic and genetic relationships of bovine natural antibodies binding keyhole limpet hemocyanin in plasma and milk. J Dairy Sci 98:2746–52

34. Elsohaby I, McClure JT, Waite LA, Cameron M, Heider LC, Keefe GP (2019) Using serum and plasma samples to assess failure of transfer of passive immunity in dairy calves. J Dairy Sci 102:567–577

35. Lutz HU, Miescher S (2008) Natural antibodies in health and disease: an overview of the first international workshop on natural antibodies in health and disease. Autoimmun Rev 7:405–9

36. McGrath B a., Fox PF, McSweeney PLH, Kelly AL (2016) Composition and properties of bovine colostrum: a review. Dairy Sci Technol 96:133–158

37. Stelwagen K, Carpenter E, Haigh B, Hodgkinson a., Wheeler TT (2008) Immune components of bovine colostrum and milk. J Anim Sci 87:3–9

38. Alipour MJ, Jalanka J, Pessa-Morikawa T, Kokkonen T, Satokari R, Hynönen U, Iivanainen A, Niku M (2018) The composition of the perinatal intestinal microbiota in cattle. Sci Rep 8:10437

39. Morrill KM, Conrad E, Lago A, Campbell J, Quigley J, Tyler H (2012) Nationwide evaluation of quality and composition of colostrum on dairy farms in the United States. J Dairy Sci 95:3997–4005

40. Reschke C, Schelling E, Michel A, Remy-Wohlfender F, Meylan M (2017) Factors Associated with Colostrum Quality and Effects on Serum Gamma Globulin Concentrations of Calves in Swiss Dairy Herds. J Vet Intern Med 31:1563–1571

41. Genc M, Coban O (2017) Effect of some environmental factors on colostrum quality and passive immunity in brown swiss and holstein cattle. Isr J Vet Med 72:28–34

42. Moore M, Tyler JW, Chigerwe M, Dawes ME, Middleton JR (2005) Effect of delayed colostrum collection on colostral IgG concentration in dairy cows. J Am Vet Med Assoc 226:1375–7

43. Godden S (2008) Colostrum management for dairy calves. Vet Clin North Am Food Anim Pract 24:19–39

44. Murphy JM, Hagey J V., Chigerwe M (2014) Comparison of serum immunoglobulin G half-life in dairy calves fed colostrum, colostrum replacer or administered with intravenous bovine plasma. Vet Immunol Immunopathol 158:233–7

45. Banks KL (1982) Host defense in the newborn animal. J Am Vet Med Assoc 181:1053–6

46. Buczinski S, Vandeweerd JM (2016) Diagnostic accuracy of refractometry for assessing bovine colostrum quality: A systematic review and meta-analysis. J Dairy Sci 99:7381–7394

47. Faber SN, Faber NE, McCauley TC, Ax RL (2005) Case study: Effects of Colostrum Ingestion on Lactational Performance. Prof Anim Sci 21:420–425

48. DeNise SK, Robison JD, Stott GH, Armstrong D V (1989) Effects of passive immunity on subsequent production in dairy heifers. J Dairy Sci 72:552–4

49. Govignon-Gion A, Dassonneville R, Baloche G, Ducrocq V (2016) Multiple trait genetic evaluation of clinical mastitis in three dairy cattle breeds. Animal 10:558–565

50. Carlén E, Strandberg E, Roth A (2004) Genetic Parameters for Clinical Mastitis, Somatic Cell Score, and Production in the First Three Lactations of Swedish Holstein Cows. J Dairy Sci 87:3062–3070

51. Ploegaert TCW (2010) Parameters for Natural Resistance in Bovine Milk.

52. Furman-Fratczak K, Rzasa A, Stefaniak T (2011) The influence of colostral immunoglobulin concentration in heifer calves’ serum on their health and growth. J Dairy Sci 94:5536–5543

53. Huang Z, Wang J, Xu X, et al (2019) Antibody neutralization of microbiota-derived circulating peptidoglycan dampens inflammation and ameliorates autoimmunity. Nat Microbiol 4:766–773

